# Echoes of 1816: Microbial Footprints in Heritage Artifacts From Argentina’s Museum of Independence

**DOI:** 10.1101/2025.03.28.645971

**Authors:** Daniel Gonzalo Alonso-Reyes, Fátima Silvina Galván, Natalia Noelia Alvarado, María Cecilia D’Arpino, Luciano José Martinez, Hernán José Esquivel, Cecilia Aymara Gallardo, María Julia Silva Manco, Virginia Helena Albarracín

## Abstract

Historical artifacts preserved in museums are invaluable cultural treasures, yet they are often vulnerable to biodeterioration caused by microbial colonization. This study presents the first comprehensive investigation of microbial communities inhabiting heritage artifacts from Argentina’s Museum of Independence. By integrating advanced microscopy with phenotypic and genomic characterization, we analyzed samples collected from wooden objects, textiles, architectural elements, and exterior walls. Scanning electron microscopy revealed diverse and well-structured biofilms, with intricate three-dimensional arrangements embedded in extracellular polymeric substances. A total of 49 bacterial strains were isolated and identified via VITEK MALDI-TOF mass spectrometry, with a predominance of Gram-positive genera such as *Bacillus, Micrococcus*, and *Kocuria*. Remarkably, the 19th-century albumen print photograph emerged as the most biodiverse artifact, yielding 21 distinct strains, including *Streptomyces, Oceanobacillus, and thermophilic Caldibacillus thermoamylovorans*. The protein-rich and halophilic environment of the albumen layer likely facilitated microbial colonization and persistence. Pseudomonas species were exclusively associated with this photographic substrate, further underscoring its niche specificity. Human-associated taxa like *Staphylococcus epidermidis* and *Staphylococcus equorum* were prevalent in high-contact areas, while exterior surfaces displayed distinct microbial signatures, including potential pathogens linked to environmental exposure. These findings reveal a complex and artifact-specific microbial landscape, emphasizing the need for tailored, bio-informed conservation strategies. This work advances the field of heritage microbiology and supports efforts to safeguard culturally significant objects through a deeper understanding of their microbial ecosystems.

## INTRODUCTION

The National Museum “Casa Histórica de la Independencia” (CHM), located in San Miguel de Tucumán, Argentina, holds a vast and culturally significant historical collection from the 19^th^ and 20^th^ centuries. The CHM is one of the most visited historical museums in the country (Chatruc, 2021) and is renowned as the site where the independence of the United Provinces of the Río de la Plata was declared on July 9, 1816 — a date later recognized as Argentine Independence Day.

The colonial mansion housing the Museum was built in 1760 and has undergone modifications over time (Marinsalda, 2015). While the original building was demolished in 1874, its 20^th^-century reconstruction has solidified its status as a national symbol of unity. The "Salón de la Jura" or Hall of the Oath, where the Declaration of Independence was redacted, remains in its original state. Apart from historical documents and furniture, the Museum conserves an extensive book library and a photographic archive. Among the highlights is Provincia de Tucumán by Arsenio Granillo, a 213-page book (Granillo, 1872) commissioned by Governor Federico Helguera, features descriptive articles and news about Tucumán, accompanied by original albumen print photographs depicting the city and its sugar mills. Notably, the book includes a unique photograph of the CHM’s original façade, considered an invaluable contribution to Argentine documentary photography and essential for the building’s restoration in 1943.

Museums serve a fundamental role in the conservation of cultural heritage (CH), safeguarding irreplaceable artifacts that embody humanity’s identity, memory, and history. Preventive conservation is vital because cultural assets are inherently vulnerable to deterioration processes that threaten their integrity (Gutarowska et al., 2014; Grabek-Lejko et al., 2017; Negi and Sarethy, 2019).

Microbiological contamination is a common issue, as many artifacts are of organic origin and prone to colonization by bacteria and fungi. These microorganisms thrive in museum environments, adapting their versatile metabolisms to withstand extreme conditions, including fluctuations in temperature, humidity, and pH. Indeed, microbes may have been present in the original piece since its first production, as invisible part of the materials used to craft the artifact. Microbes then settled into specific ecological niches, often remaining undetected until visible biofilm formation, discoloration, or structural weakening became apparent. Their activity can cause significant damage to textiles, wood, paper, and other organic materials, manifesting as discoloration, structural weakening, odor, and cracking (Skóra et al., 2015; Grabek-Lejko et al., 2017; Negi and Sarethy, 2019). The consequences of microbial growth extend beyond artifact damage, potentially diminishing monetary value, and bringing about costly decontamination procedures (Sterflinger and Piñar, 2013). Despite these challenges, the cultural and historical value of many CH objects is immeasurable and cannot be quantified solely in monetary terms.

Biodeterioration is defined as “any undesirable change in a material brought about by the vital activities of organisms”(Perito and Cavalieri, 2018). Although abiotic processes were historically viewed as the primary cause of CH decay, the role of microorganisms has gained recognition only recently. While biodeterioration cannot be entirely halted, advances in microbial ecology and technological interventions offer strategies to mitigate its effects. A deeper understanding of microbial communities on CH objects is critical for identifying their deteriorative mechanisms and implementing effective control measures (Perito and Cavalieri, 2018). These aspects have become an important research field for both, conservators and microbiologists (Adamiak et al., 2018). This issue also raises occupational health concerns, as museum staff may be exposed to harmful biological agents during the handling of contaminated materials, such as paper, wood, and textiles (Borrego et al., 2010; Gutarowska et al., 2014). Indeed, many studies indicated that conservators, librarians, and archivists face risks related to microbial exposure (Zielińska-Jankiewicz et al., 2008; Wiszniewska et al., 2009; Karbowska-Berent et al., 2011).

Traditional methods of investigating microorganisms on CH objects have primarily been culture-dependent, offering insights into their deteriorative roles. Recently, amplicon sequencing has emerged as a powerful yet resource-intensive tool for microbial identification (Kraková et al., 2012a, 2018). An alternative approach combines the automated VITEK system for rapid biochemical identification (Książczyk et al., 2016), with whole-genome sequencing (WGS) for functional profiling of key strains. Scanning electron microscopy (SEM) has also proven effective in biodeterioration studies, providing visual insights into microbial activity on CH surfaces (Díaz-Herráiz, 2015; Puškárová et al., 2016; Elhagrassy, 2018; Liu et al., 2018).

To better understand the role of microbial communities in the biodeterioration of cultural heritage artifacts, we employed a multidisciplinary approach integrating advanced microscopy, culture-dependent techniques, and genomic analyses. Scanning electron microscopy (SEM) was chosen to visualize microbial colonization and biofilm structures on historical surfaces at high resolution, while culture-based methods allowed us to isolate and characterize viable bacterial strains. The application of MALDI-TOF mass spectrometry enabled rapid and accurate taxonomic identification, and whole-genome sequencing provided insights into the functional potential of selected isolates, particularly their capacity for biodegradation and resistance to environmental stressors. This multidisciplinary approach allows us to analyze the microbial communities in detail, revealing their influence on the artifacts’ preservation and contributing to informed conservation practices.

## 2. MATERIALS AND METHODS

### 2.1. Selection of heritage artifacts of the Museum

The CHM is located in San Miguel de Tucumán, the capital of Tucumán Province, Argentina at a pedestrian zone with significant foot traffic. (https://sketchfab.com/3d-models/museo-nacional-casa-historica-5b37f88491c0456bbd15fe87f4b8b171). The Casa Histórica is a preserved colonial-style building and functions as a museum, offering guided tours and exhibitions related to Argentina’s independence and the revolutionary era. The museum features four courtyards, a cistern, and native trees from the region. In 1941, the house was declared a National Historic Monument.

CHM currently holds over 700 items in its collection. This includes the original Act of the Declaration of Independence and various other pieces such as period furniture, weapons, personal items, dishware, religious artifacts, paintings, portraits, coins, and commemorative plaques and medals from the 18^th^ and 19^th^ centuries. The museum’s library contains classic texts, recent historiographical debates, museum studies, conservation, restoration, literature, journals and early provincial newspapers. With over 500 volumes, the collection covers diverse topics such as ecclesiastical history, the Wars of Independence, civil wars, art, the sugar industry, economics, travel narratives, Jesuit chronicles, and journalism, with a focus on the province of Tucumán. Before sampling, an extensive survey and photographic documentation were conducted to capture the condition of cultural heritage items affected by different types of deterioration. This preliminary assessment informed the selection of specific sampling locations within the building and on key artifacts of cultural significance.

The initial phase of sampling took place in August 2019. Samples were taken from the museum’s exterior, including facade elements like the front door and painted walls (Fig. 1A), which are subjected to annual repainting, weathering, and urban pollution; they were thus considered a control zone related to outdoor urban microbiome (OUM). The front was built as part of the last reconstruction in 1943 including the wooden door initially coated with flax oil. Within the museum, samples were taken from the window and furniture in the “Salón de la Jura”. Additional historical artifacts (points 2.2 and 2.3) that required preservation evaluations were also sampled (Table 1, Fig. 1). All these samples were considered as specimens of the indoors museum microbiome (IMM).

**Figure 1.** Sampled locations of the CHM. **A.** Facade. **B.** “Salon de la Jura”. **C-E**. Chair and table from the “Salón de la Jura”. **F-G.** Window Salon de la Jura. **H-I.** Alberdi’s garments. **J-L.** Albumen print book. **M.** Washing trough. **N.** Main door.

**Table 1.**
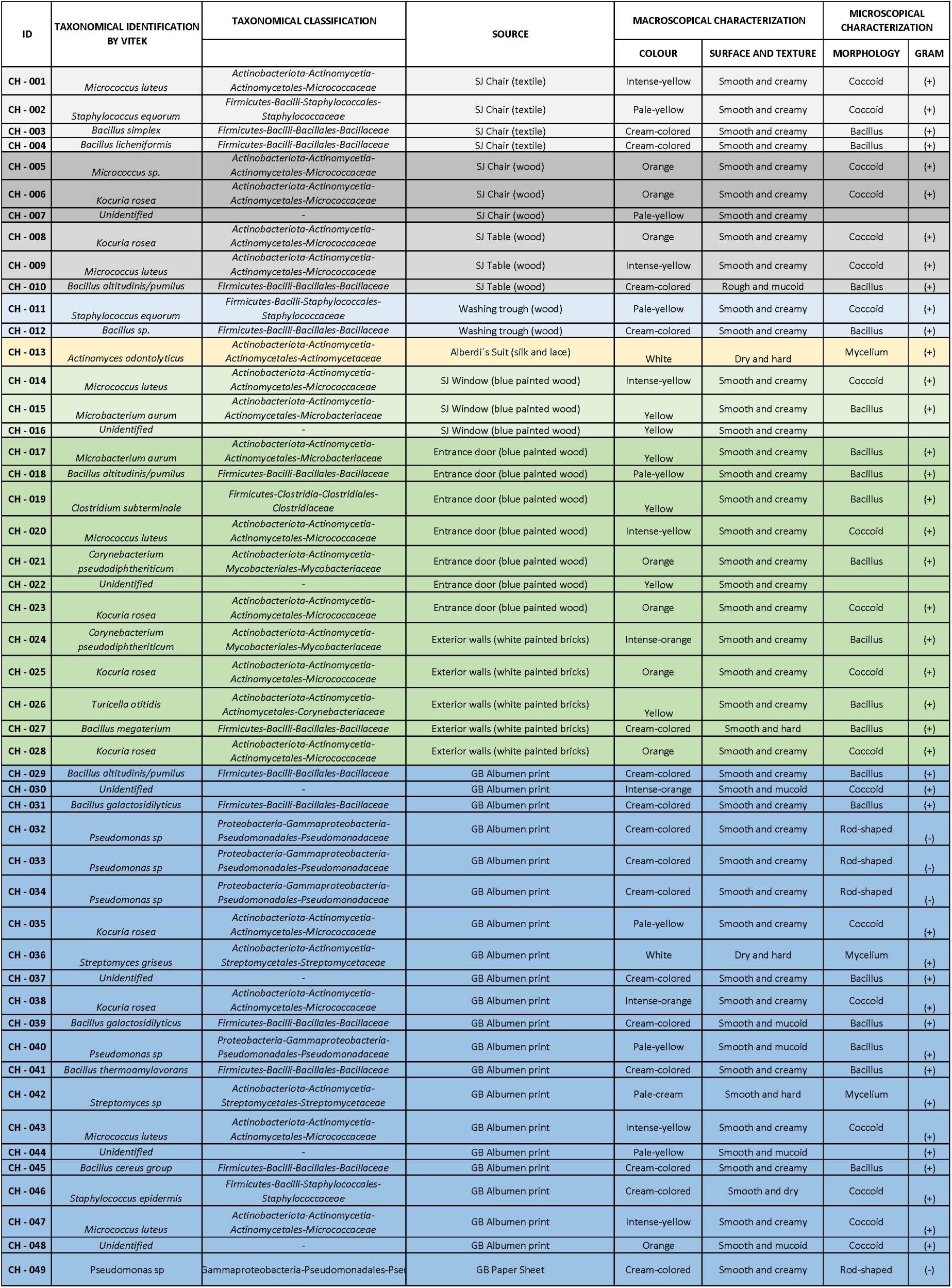
Taxonomical, macroscopical and microscopical details for most of the 49 strains isolated from the CHM.

### 2.2. Salón de la Jura

The Salón de la Jura or Hall of the Oath (SJ) is a central room within the CHM (Fig. 1B). This site holds historical significance as it is where the Congress of Tucumán proclaimed Argentina’s independence from Spanish rule on July 9, 1816. The hall is renowned for its role as the setting of intense debates and the eventual drafting of the "Acta de la Independencia," marking the foundation of the Argentine nation.SJ is the only room of the house that has been conserved intact since then. The room features period furnishings and artifacts that transport visitors back to the momentous occasion when representatives from the United Provinces of the Río de la Plata gathered to declare sovereignty. Samples for our study were collected from one chair where congressmen sat and from the table where the Declaration of Independence was signed; this furniture remains in its original state, dating back to 1810.The wooden frame from the armrest and backrest and the velvet-covered seat of one of the chairs were sampled to assess the microbial communities present on these distinct materials (Fig. 1C-D). The table’s wooden surface was swabbed in multiple locations, especially in the areas less exposed to regular cleaning, to try to sample the original microbiome in the furniture (Fig. 1E). Additionally, we took samples from biodeteriored zones of one of the windows of the room, subjected to infestation by woodworm eggs (1F-G).

### 2.3. Child Suit from Juan Bautista Alberdi, the father of Argentinean Constitution

Juan Bautista Alberdi (1810-1884) was a distinguished statesman, diplomat, and writer who played a pivotal role in shaping Argentina’s political landscape. Born in San Miguel de Tucumán, he was known for his influential work in drafting the Argentine Constitution of 1853. Thus, Alberdi’s legacy as a thinker and leader has left a lasting impact on the country’s history and development.

The Museum holds a captivating collection of artifacts attributed to Alberdi, including an item that offers a rare and personal look at 19^th^-century life: a childhood suit made from natural red silk (Catalog Number: CH321; 53 cm x 29 cm; sewing technique manufactured in Argentina [Fig. 1H-I]), This garment reflects the fashion of adult male clothing of the period, characterized by its fabric, design, and color. The suit comprises two main pieces: a jacket featuring a draped front center with a faux belt detail and a pleated back skirt; the trousers are adorned with decorative lace and bows at the side hems. These garments serve as a tangible connection to Alberdi’s early life and provide a unique perspective on the sartorial and social customs of the era. Its historical significance is enriched by its association with one of Argentina’s key historical figures, illustrating how clothing can encapsulate cultural identity and societal norms. The Museum has employed preventive conservation strategies, including controlled humidity, light, and temperature, to maintain the suit’s condition and ensure its longevity. https://sketchfab.com/3d-models/traje-de-juan-baustista-alberdi-037b9c3badd54d21b330a3d25e2b91d3

### 2.4. An iconic photograph on the Historical Book *Provincia de Tucumán*

The second phase of sampling in December 2021 targeted a key artifact from the CHM: the 1872 book *Provincia de Tucumán. Serie de artículos descriptivos y noticiosos*, authored by Dr. Arsenio Granillo (Fig. 1J-L). Commissioned by Governor Federico Helguera, this work is historically notable for its pioneering inclusion of albumen prints captured by Italian-born photographer Ángel Paganelli. These early photographs, which include a depiction of the Casa Histórica façade, provide rare insights into Tucumán’s 19^th^-century landscapes and urban spaces.

Stored in a metallic safe case to preserve its condition, this book has survived with minimal light exposure, thermal fluctuations, and limited handling. Of the estimated 300 original copies printed, only about 100 featured the albumen prints, and few remain today—primarily in the Casa Histórica and private collections.

The albumen prints stand out both for their historical value and the meticulous craftsmanship involved in their creation. Paganelli used large glass negatives (approximately 15 x 12 cm) to produce contact prints on albumen-coated paper. This egg-white and sodium chloride mixture served as a binder for silver salts, enabling high-quality image transfer. The labor-intensive process involved exposure to sunlight under negatives, toning, fixing, and washing the images (Reilly, 1980) before affixing them to book pages by hand. Each of the 100 photograph-included books contained 21 images, totaling over 2,100 individually prepared prints—a remarkable testament to 19th-century photographic artistry.

Due to their production method, we considered albumen prints as halophilic microenvironments, which may influence microbial colonization. The unique environmental conditions within the book highlight the delicate balance required for preserving such historical artifacts while exploring their microbiological significance.

### 2.5. Sampling procedures for direct material observation and bacterial isolation

To conduct the sampling, a non-invasive approach was employed, using precision tools to extract material samples without compromising the structural integrity of the pieces. Micro-sampling techniques, supported by visual documentation and environmental controls, were used to assess the physical condition of the albumen prints and the rest of the elements identify potential areas of degradation, such as yellowing, flaking, or damage from previous handling.

For the direct observation of microorganisms on heritage objects, freshly-open 3M™ adhesive tape was used (Michaelsen et al., 2009). Briefly, tape was cut into strips (2-3 cm), subjected to 15-min sterilization under UV-C. Subsequently, the strips were gently adhered to the different surfaces and then transferred to 2 ml tubes containing Karnovsky’s fixative solution (2.66% w/v paraformaldehyde and 1.66% w/v glutaraldehyde in 0.1 M phosphate buffer, pH 7.2), for transportation to the lab. The tape sampling method was not used for the book of Granillo, to avoid any undesired consequences over its conservation.

For isolation of individual bacterial strains, sample collection was performed with dry and sterilized cotton swabs. The swab was passed over the entire area of visibly damaged material; each swab covered approximately a 10 cm² area to standardize sample collection. Immediately after sampling, the swabs were used to inoculate plates with Luria-Bertani (LB) agar media. Parallel agar plates were supplemented with nalidixic acid (10 µg mL-1) and cycloheximide (10 µg mL-1) to inhibit the growth of Gram-negative bacteria and fungi, respectively (Rasuk et al., 2017). This non-invasive method is considered essential for the study of collections in museum environments (Michaelsen et al., 2009; Piñar et al., 2015).Once the colonies appeared, they were collected with a sterile loop for purification and subsequent streaking allows generating axenic cultures. We stored the cells in glycerol 20% at -20°C. Subcultures were performed in LB broth for morphological and molecular identification analysis. From the pure cultures of the bacterial isolates, macroscopic practices were carried out to determine the structure of the colonies and their pigmentation, and microscopic practices (optical and electronic) of individual cells to establish shapes, sizes, and spatial arrangement. Additionally, the Gram reactions of all isolates were recorded.

All isolated strains in this work are kept and curated in the CEBAC collection (Cepario de Bacterias Ambientales del CIME), Catalogue Urban Microbiome Strains, under the corresponding codes indicated in Table 1.

### 2.6. Scanning Electron Microscopy (SEM)

For SEM studies, the adhesive tapes or pure axenic cultures of isolated strains were fixed in a mixture of aldehydes (Karnovsky fixative) overnight at 4°C, and were prepared following the processing method already previously described in Alonso-Reyes et al (2021). During post-fixation, samples were washed with 0.1 M phosphate buffer twice and dehydrated in a graded concentration series of ethanol (50, 70, 90, and 100%), following acetone 100%. This was followed by critical point drying (Denton Vacuum model DCP-1). The samples were then mounted on aluminum supports and subjected to gold sputtering (JEOL model JFC-1100) and examined using a Zeiss Supra 55VP scanning electron microscope (Carl Zeiss NTS GmbH, Germany), in the Electron Microscopy Core Facility (CIME-CONICET-UNT).

### 2.7. Identification of microbial strains by MALDI-TOF MS (VITEK)

MALDI-TOF analysis was performed with a bioMérieux VITEK MALDI-TOF mass spectrometer (bioMérieux, France), and the spectra were compared to the VITEK MS SARAMIS database for research use only (RUO), version 4.09, using the SuperSpectra algorithm (referred to here as MALDI-TOF MS). For MALDI-TOF MS analysis, a colony from a pure plate with 24h growth was applied to a slide and covered with 1 µl of matrix solution (α-cyano-4-hydroxycinnamic acid) and dried completely before analysis. The identification was considered definitive when the probability provided was greater than 70%. This service was provided by Laboratorios CIBIC in Rosario, Argentina.

### 2.8. DNA extraction

DNA from eight isolated strains was obtained using a commercial extraction kit (DNA Puriprep B-kit for bacteria, Inbio Highway, Argentina); following the manufacturer’s instructions. For Gram-positive bacteria, the pellet was suspended in 162 µl of BRB buffer and 18 µl of lysozyme, and then incubated for 30 min at 37°C. For gram-negative bacteria, the use of lysozyme was omitted. Next, 200 µl of BT buffer and 20 ul of proteinase K were added to each sample. It was incubated for 30 min at 56°C. After, RNaseA (10 mg/ml), was added and incubated for 30 min at 37°C. Then the lysis buffer (200 µl) was added, vortex pulsed for 10 sec and the homogenate was incubated for 10 min at 56°C. Then, 200 µl of 100% ethanol was added to each sample, shaken by inversion, and transferred to a microcolumn. It was centrifuged for 1 min at 12,000 g, and the filtrate was separated. Then, two column washes were performed each with 500 µl of Blav1 and Blav2 buffer respectively. Finally, the column was dried by centrifuging for 3 min at 12,000 g. DNA elution was carried out with 60 µl of pH 9 elution buffer, equilibrated at 70°C. DNA concentration and quality were measured using a µDrop plate (Thermo Scientific).

### 2.9. Genome sequencing, assembly, and analysis

Whole genome sequencing service was performed by Novogene UK services (https://www.novogene.com/amea-en/) using the IlluminaNovaseq platform with pair-end 150 strategy. Quality control of the reads was also performed by Novogene. The Illumina reads were assembled using SPAdes assembler v. 3.15.4 (Bankevich et al., 2012), with -- careful and --cov-cutoff auto options. The resulting eight assemblies were uploaded to the NCBI database under the Bioproject accession number PRJNA1233209, and are also available at Zenodo (https://doi.org/10.5281/zenodo.15169295). The functional annotation was performed using Arche v. 1.0.1 (Alonso-Reyes and Albarracín, 2022) pipeline with -- kegg option, and the resultant KEGG identifiers (KO) were mapped through the KEGG mapper reconstruct tool (https://www.genome.jp/kegg/mapper/reconstruct.html). The genomic potential for the synthesis of secondary metabolites was elucidated through antiSMASH v. 7.1.0 (Blin et al., 2023). To estimate the taxonomic identity of each strain at genus level, 16s-rDNA and DNA gyrase B genes were retrieved from the genomes using barrnap and blast tools, and then blasted against the NCBI NR database. For complete taxa identification, average nucleotide identity (ANI) analysis was performed using pyani (Pritchard et al., 2016), against genus-specific genomes deposited in NCBI database.

## 3. RESULTS

### 3.1. SEM analysis of microbial communities on heritage objects and façade surfaces

Biofilms on heritage surfaces including furniture, wooden objects (chair, table, window and washing trough) and textile garments were analyzed using scanning electron microscopy (Fig. 2). The freshly painted building façade, entrance door and exterior walls, were considered as control samples of the OUM for comparison with the IMM. In all cases, the adhesive tape sampling method proved to be a useful, simple, and non-destructive technique for collecting microbial communities for immediate imaging analysis. Interestingly, the surfaces of objects within and outside the museum were highly colonized with microbes forming well-structured biofilms.

**Figure 2.** Biofilms on heritage surfaces including furniture, wooden objects (chair, table, window and washing trough) and textile garments. **A.** Chair. **B.** Table. **C-D.** Washing trough. **G-H.** Garments. **I.** Window. **J.** Entrance door.

Most observed biofilms exhibited three-dimensional cellular organization and were surrounded by substantial amounts of extracellular material. For instance, the wooden chair showed spindle or hazelnut-shaped bacterial cells embedded in an abundant exopolysaccharide (EPS) matrix (Fig. 2A). The wooden table displayed a homogeneous bacterial biofilm with coccoid units connected by thin, short filaments (Fig. 2B). In the wooden washing trough, compact coconut-shaped biofilms were observed attached to wooden rests (Fig. 2C), while forming an interconnected by extracellular filaments or vesicles. Other areas revealed a filamentous bacterial mantle (Fig. 2D) likely belonging to the genus *Streptomyces* or related Actinobacteria. On garments, microbial filaments were much scarce (Fig. 2G-2H); they were adhered tightly to deteriorate fibers of the clothing, with fibrous materials and deposits visible at higher magnifications. In this sample, diverse fungi spores were also found. In the window sample, we observed a biofilm formed by heterogeneous microbial community dominated by large cocci and compacted bacilli, with minimal extracellular material facilitating cell interactions (Fig. 2I).

The façade also showed high colonization of the surfaces by biofilms; the entrance door exhibited a thick biofilm structure composed of cocci, dividing bacilli, larger rough-surfaced bacilli, and filamentous bacteria, all forming strongly interconnected aggregates (Fig. 2J). The exterior walls demonstrated consolidated coccoid biofilms held together by EPS, likely offering protection against environmental factors, while other regions showed sparse adherence of bacilli in simple distributions.

### 3.2. VITEK-MS-based Identification of Isolated Bacteria

A total of 49 bacterial strains were isolated from various samples (Table 1, Fig. 3-4), with 12 strains attributed to the OUM and 37 strains linked to the IMM. Each sampled niche exhibited a distinct profile of bacterial taxa, reflecting the contrasting conditions of outdoor and indoor environments. Moreover, each artifact sampled from within the Casa Histórica de la Independencia can be regarded as an isolated microbial island, shaped by its unique environmental conditions, material composition, age, usage and historical context. The microbial communities inhabiting these artifacts form a complex and dynamic microecosystem, where each niche harbors a distinctive collection of bacteria specifically adapted to its substrate (Fig. 3). Gram-positive bacteria dominated the isolates, as expected given the use of selective isolation media supplemented with antibiotics to exclude fungi and gram-negative bacteria. Nevertheless, five strains of *Pseudomonas* were successfully isolated under these same conditions from the Granillo Book—one from the paper and four from the photograph itself. Indeed, the most prolific source of isolates was the albumen photograph, from which 21 distinct strains were recovered, predominantly from the genera *Bacillus, Oceanobacillus, and Streptomyces*. In contrast, the Alberdís garment gave a unique isolate: *Actinomyces odontolyticus*. Morphological characteristics of each strain (texture, color, spore production) were recorded to differentiate isolates (Table 1).

**Figure 3.** Illustration of the CHM and sampling sites. Several taxonomical groups assigned to the isolates through VITEK are represented by symbols.

Four strains were isolated from the textile upholstery of a chair (*Micrococcus luteus, Staphylococcus equorum, Bacillus licheniformis and Bacillus simplex*) while three strains were found on the wooden frame of the same chair (*Micrococcus* sp., *Kocuria* sp. and one unidentified strain). Additional 15 isolates were recovered from other wooden elements, including the SJ table (*Kocuria rosea, Micrococcus luteus, Bacillus altitudinis/pumilus)*, the SJ window frame (*Micrococcus luteus, Microbacterium aurum* and one unidentified strain), the museum’s entrance door (*Microbacterium aurum*, *Bacillus altitudinis/pumilus*, *Clostridium subterminale*, *Micrococcus luteus*, *Corynebacterium pseudodiphtheriticum, Kocuria rosea* and one unidentified strain) and a washing trough (*Staphylococcus equorum* and *Bacillus* sp.) where meat of slaughter animals was prepared. Finally, five strains were isolated from the front wall of the building made of white painted bricks (*Corynebacterium pseudodiphtheriticum, Kocuria rosea, Turicella otitidis, Bacillus megaterium, Kocuria rosea*).

### 3.3. Genome-based taxonomical identification

The genomes of eight representative strains were sequenced and analyzed for functional characterization. Three of them correspond to bacterial isolates from the front and will be considered as representatives of the OUM while the rest are considered representatives of the IMM (Table 2). The sequencing process and quality control yielded approximately 100 million reads and 15 Gb of data, with an average of 12 million reads and 1.88 Gb per genome. Assembly resulted in fewer than 115 contigs for all strains, except for the CH-041 strain, which had 200 contigs. The G + C content varied between strains, ranging from ∼35% in CH-031 to ∼72% in CH-021. The ARCHE pipeline, employing GeneMarkS-2 for gene prediction, achieved high annotation rates for protein-coding sequences (CDS). The lowest annotation rate was observed in CH-026, with ∼85% of CDS annotated, while CH-036 reached approximately 97% annotation.

**Table 2.**
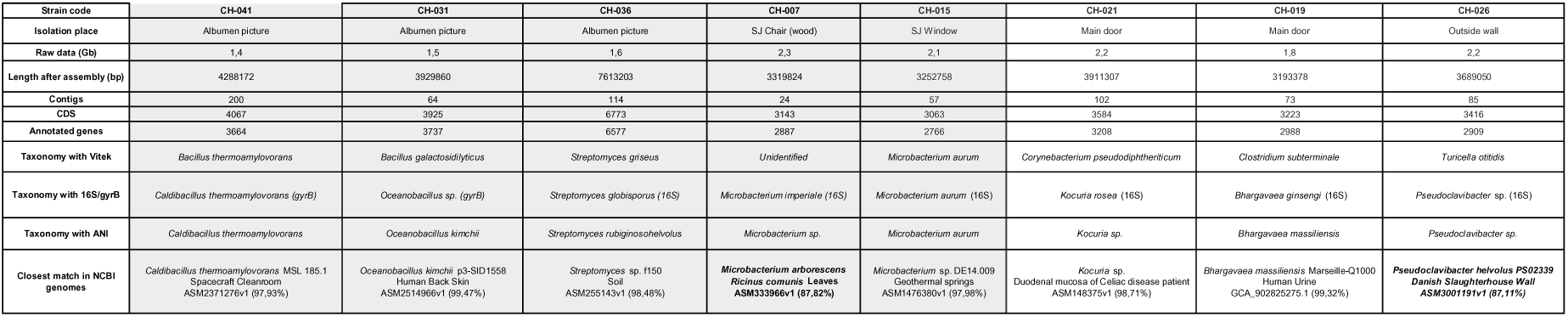
Data of the sequenced isolates, including the isolation place, sequencing results, de novo assembly, annotation, and taxonomical identification by several methods. In grey, are the Indoors Museum Microbiome (IMM) isolated strains, in white the Outdoors Urban Microbiome (OMM) ones.

Species identification, initially conducted using the automated VITEK system (Książczyk et al., 2016) was supplemented with 16S-rDNA and DNA gyrase B marker genes (MGs) against NCBI databases. The gyrB gene was used when the 16S-rDNA gene was incomplete or truncated. ANI analyses were performed using genus-specific genomes from the NCBI database for more precise identification (Table 2).

VITEK identification results for most strains differed from those obtained through MGs. Only strains CH-041 and CH-015 maintained the same species classification by both methods. CH-036 was identified as belonging to the genus *Streptomyces* by VITEK, MGs, and ANI, but the species differed between methods. VITEK accurately classified CH-031 at the family level (Bacillaceae), but MGs identified it at the genus level as *Bacillus*, whereas ANI suggested *Oceanobacillus*. CH-021 and CH-026 only matched at the class level (Actinomycetes) by both methods, while CH-019 matched at the phylum level (Bacillota).

CH-007 could not be classified by VITEK but was identified as *Microbacterium* by MGs and ANI. Significant differences were evident in species-level classifications; MGs and ANI agreed on the species for CH-041 (*Caldibacillus thermoamylovorans*) and CH-015 (*Microbacterium aurum*). In CH-031, where MGs did not yield a species classification, ANI clustered it with *Oceanobacillus kimchi* due to >95% similarity to a reference genome in the NCBI database (Richter and Rosselló-Móra, 2009). Notably, CH-007, and CH-026 did not align with any reference genome above the 95% ANI threshold, suggesting they might belong to novel species.

Pairwise comparisons of the CHM genomes with public database genomes revealed the closest matches. Most strains shared >97% similarity, except for CH-026 (87.11%) and CH-007 (87.82%). For example, CH-031 had a 99.47% sequence identity with *Oceanobacillus kimchii* p3-SID1558 (Accession No. <u>NZ_JALXUQ000000000</u>), isolated from human back skin (Hannah et al., 2023). CH-019 showed a 99.32% identity with *Bhargavaea massiliensis* Marseille-Q1000 (Accession No. <u>NZ_JAHBCJ000000000</u>), from a urine sample of patients with urinary tract infections (Olowo-okere et al., 2021). CH-021 matched with *Kocuria* sp. CD08_4 (Accession No. <u>NZ_LQBK00000000</u>) at 98.71%, an isolate from the duodenal mucosa of a celiac disease patient, which carries genes linked to virulence and oxidative stress resistance (Munish et al., 2017). As expected, CH-036 matched a *Streptomyces* strain with 98.48% of similarity, *Streptomyces* sp. f150, which was isolated from soil at Florida, USA (Accession No. <u>NTHG00000000.1</u>). CH-015 showed a 97.98% similarity with the metagenome-assembled genome (MAG) of *Microbacterium* sp. DE14.009 (Accession No. <u>JACXUU000000000.1</u>) from geothermal springs in Greece. CH-041’s genome matched *Caldibacillus thermoamylovorans* MSL 185.1 (Accession No. <u>NZ_JAMAXM000000000</u>), a thermophile isolated from a spacecraft environment, capable of enduring heat-shock at 80°C for 15 minutes (Tran et al., 2023). In contrast, CH-007 and CH-026 were related at a distance to *Microbacterium arborescens* RCB1 (87.82% similarity; Accession No. <u>QNUW00000000</u>) and *Pseudoclavibacter helvolus* PS02339 (87.11% similarity; Accession No.<u>NZ_JASCYB000000000</u>), respectively, and do not meet the 95% threshold for known species. Genetic comparisons indicated approximately 12% divergence from their closest known relatives in the NCBI database, indicating that both may be novel species.

### 3.4. Functional Annotation of Genomes: Insights into Biodeterioration and Environmental Resilience

The functional annotation carried out using ARCHE led to useful information regarding the genomic potential of the strains (Fig. 5.A-E, Supplementary Tables S1-15). Genes involved in resistances to antibiotics, osmotic stress, and heavy metals were reported from the annotation. Furthermore, relevant genes related to the biodeterioration of the cultural heritage were found, such as those required for the degradation of wood and paper, and the production of pigments. *Streptomyces* sp. CH-036 revealed the highest numbers of genes in all the systems mentioned before; except for heavy metal resistance (the numbers are still relatively high). Moreover, CH-036 reported more than twice as many genes related to biodeterioration (Fig. 5.D-E) as the rest of the strains studied. Particularly, those genes related to the degradation of cellulose and production of pigments, which are potentially harmful to paper documents, were highly abundant in CH-036. *Kocuria* sp. CH-021 strain displayed a high potential to resist both the osmotic and heavy metal stresses. A relatively elevated number of cation transporters were annotated in CH-021’s genome, and genes related to copper and chromate homeostasis were also noticeable. Genes for the degradation of cellulose and hemicellulose are also present, together with the capability to produce some pigments. On the other hand, both *Ocenobacillus* sp. CH-031 and *Caldibacillus* sp. CH-041 had a reduced biodeteriorative potential; a tiny group of genes devoted to pigment production and degradation of wood and paper were found (none of the latter are related to cellulose degradation). Finally, CH-041 showed a clear heavy metal resistance profile considering the coding-sequences found in the genome.

**Figure 4.** Subcultures performed in LB broth for some CHM strains, complemented by gram staining and SEM microscopy. **A.** CH-005. **B.** CH-003. **C.** CH-010 **D.** CH-011. **E.** CH-013 **F.** CH-015. **G.** CH-021 **H.** CH-026.

**Figure 5.** The genomic potential of eight sequenced strains unveiled by functional annotation. **A-E.** In barplots are depicted the abundances for those genes involved in resistances to antibiotics, osmotic stress, heavy metals and those required for the degradation of wood and paper, and the production of pigments. **F.** Potential for the synthesis of secondary metabolites, and degradation of xenobiotics.

Further functional capabilities of relevance were studied, such as the potential for the synthesis of secondary metabolites, and degradation of xenobiotics (Fig. 5.F). The results of the screening with antiSMASH revealed *Streptomyces* sp. CH-036 as the most diverse of the strains regarding to the hosting of biosynthetic gene clusters (BGCs) devoted to the production of secondary metabolites. Among the putative products of these BCGs are lanthipeptides, and ε-poly-L-lysines, both compounds with reported antimicrobial properties (Repka et al., 2017; Wang et al., 2021). Other compounds reported from CH-036 genome were desferrioxamin (Holden and Nair, 2019), and isorenieratene, which is a light-induced yellow pigment (Takano et al., 2005). On the other hand, KEGG identifiers retrieved from the annotation of *Kocuria* sp. CH-021 genome revealed several complete pathways for xenobiotic degradation, including those for the degradation of benzoate, phenylacetate, xylene, and toluene.

## 4. DISCUSSION

### 4.1. Microbial Niches in the Museum and Their Impact on Heritage Conservation

This study revealed significant microbial colonization on historical artifacts and architectural surfaces, highlighting both the conservation challenges and the potential scientific value of these microbial communities. SEM analysis of historical artifacts showed extensive biofilm development in most samples. In some cases, like the Alberdís garment, it was observed tightly adhered microbial filaments to altered original material of the pieces, providing direct evidence of biodeterioration processes that threaten cultural heritage.

Microbial colonization in the museum environment is shaped by distinct ecological niches, with significant differences observed between the outdoor urban microbiome (OUM) and the indoor museum microbiome (IMM). The OUM, primarily represented by microbial communities on the museum’s façade, entrance door, and exterior walls, is continuously exposed to fluctuating environmental conditions such as temperature variations, UV radiation, precipitation, and air pollutants. This exposure selects for stress-resistant taxa thriving under changing conditions, including endospore-forming *Bacillus* spp., UV-resistant *Micrococcus luteus*, and environmental actinobacteria, which form resilient biofilms. For instance, *Microbacterium aurum* is known for its environmental robustness and ability to survive desiccation and nutrient limitations, while *Bacillus altitudinis/pumilus* forms endospores that protect against UV radiation and other stressors. The presence of species such as *Kocuria rosea*, known for pigment production and environmental stress resistance, further reflects the selective pressures of urban niches. Additionally, airborne contaminants and human activity in urban settings introduce anthropogenic microbes, some of which may be opportunistic pathogens (*Pseudoclavibacter* sp., *Kocuria* sp., *Bhargavaea* sp.) or biofilm producers that facilitate biodeterioration (Stackebrandt et al., 1995; Bartlett et al., 2022; Zhou et al., 2022; Ahamad et al., 2023). Furthermore, when ANI comparisons are used the genome of the strain CH-019 shares a 99,32% of similarity with *Bhargavaea massiliensis* Marseille-Q1000 isolated from a urine sample in patients with urinary tract infections (Table 2, Section 3.3). Likewise, ANI showed a 98,71% of similarity between the genomes of CH-021 and *Kocuria sp.* CD08_4 an isolate from the duodenal mucosa of a celiac disease patient. Finally, CH-017 and CH-027 were both classified by VITEK as *B. pumilus* and *B. megaterium* which have been reported as plant pathogens in other work (Bull et al., 2010; Yuan and Gao, 2015).

In contrast, the IMM comprises microbial communities inhabiting historical furniture, textiles, paper artifacts, and architectural surfaces within the controlled indoor environment of the museum. These niches are characterized by relatively stable temperature and humidity conditions, lower UV exposure, and reduced air circulation, which favors specialized microbial assemblages. Here, bacterial communities may be defined by the original organic and inorganic substrates of the artifact and the use given to the museum piece. Wooden furniture, including the chair and table, was dominated by *Bacillus pumilus*, *Micrococcus* sp., and *Kocuria rosea*, which are adapted to nutrient-limited and stable conditions. Alberdi’s suit yielded only one strain, *Actinomyces odontolyticus*, a filamentous bacterium well-suited to organic substrates like fabric fibers and related with dental infections. The albumen photograph exhibited exceptional diversity, including *Caldibacillus thermoamylovorans*, *Oceanobacillus kimchii*, and *Streptomyces rubiginosohelvolus*, showcasing the unique properties of albumen as a protein-rich substrate conducive to microbial colonization. The dominance of *Actinomyces*, *Staphylococcus*, and *Bacillus* species in the IMM suggests a strong association with human contact, dust deposition, and material degradation. Notably, *Pseudomonas* spp., exclusively isolated from the albumen photograph, may play a role in gelatin and paper biodeterioration, highlighting the niche specificity of microbial colonization. These findings emphasize the IMM’s potential as a window into historical microbial communities and their interactions with cultural heritage substrates, offering valuable insights into microbial persistence and biodeterioration. Simultaneously, they highlight the need for targeted conservation strategies to address microbial degradation while preserving the historical significance of these microbial communities.

Indeed, it was pointed out the importance of regular disinfection routines, after several *Bacillus* spp., *Staphylococcus* spp., and *Streptomyces* spp. with potentially pathogenic characteristics were reported in museum collections (Gutarowska et al., 2014). Furthermore, it has been suggested that some *Streptomyces* spp. present in museums could be noxious to the human respiratory system (Borrego et al., 2010). In our work, we found the strains CH-036 and CH-042 classified as *Streptomyces* spp. by different methods. Surprisingly, the genome of CH-036 showed a strong antibiotic resistance profile of genes (Fig. 5.A). It is thus important keep away threats to health from the CHM’s staff and visitors, especially those who are immunocompromised. Protection elements should be used in the manipulation of museum artifacts. Also, we encourage the adoption of appropriate measures to avoid contamination of the CHM’s elements with new infectious agents coming from the outside of the CHM’s. Proof of the unprotected handling of the book of albumen print is the presence of the commensal species *Staphylococcus epidermis*, related to the human skin microbiome (Villalba et al., 2004). Also, a 99.47% of similarity was found between the genomes of CH-031 and *Oceanobacillus kimchii* p3-SID1558 isolated from the human back skin (Table 2). In this case, as photographs were hand-crafted prepared and then pasted in the book, an initial inoculation of the strain from the original photographer ’skin microbiome cannot be rule out.

The interaction between OUM and IMM microbiomes is a critical yet underexplored aspect of microbial dynamics in heritage conservation. Airborne microbial transfer, visitor-mediated contamination, and ventilation systems could facilitate cross-colonization between outdoor and indoor environments. Additionally, spores and biofilm fragments from external surfaces may infiltrate indoor spaces, potentially influencing microbial succession on artifacts. Understanding these interactions is essential for designing targeted conservation strategies that mitigate microbial spread while preserving the unique ecological stability of indoor heritage collections. Further studies incorporating metagenomic and metatranscriptomic analyses could provide deeper insights into microbial connectivity and functional roles in museum environments.

### 4.2. Photographic heritage as novel source for polyextremophilic bacteria

The albumen prints in the book Provincia de Tucumán represent a unique microenvironment that may influence microbial colonization and biodeterioration processes. Albumen, composed primarily of denatured egg white proteins, creates a proteinaceous, nitrogen-rich substrate with high organic content, potentially favoring the growth of microorganisms capable of metabolizing proteins. This composition makes albumen prints particularly susceptible to colonization by proteolytic bacteria and fungi, which may exploit albumen as a carbon and nitrogen source. Moreover, the high hygroscopicity of albumen paper, resulting from both the organic matrix and the hydrophilic nature of its gelatinous binder, creates microclimatic conditions conducive to microbial proliferation. Fluctuations in relative humidity (RH) can lead to the absorption and retention of moisture, facilitating the activation of dormant microbial spores and the formation of biofilms. The presence of *Pseudomonas* spp., a known gelatin and protein degrader, exclusively in albumen-coated prints, supports this hypothesis, suggesting that this niche selects for microbial taxa with proteolytic capabilities.

Another key factor is the interaction between albumen, silver salts, and environmental pollutants. Albumen prints were historically prepared with silver nitrate, which reacts with the albumen matrix to form light-sensitive silver salts. While silver ions possess antimicrobial properties, over time, their oxidized and degraded forms may lose effectiveness, allowing microbial colonization in aged prints. Additionally, sulfur-containing air pollutants (e.g., hydrogen sulfide) could interact with residual silver, forming silver sulfide (Ag S), which is less antimicrobial and contributes to the well-known yellowing and fading of albumen photographs. We hypothesize that the combination of albumen’s organic composition, moisture retention, and silver-based chemical interactions creates a selective microbial niche that differs from other paper-based heritage objects. The albumen photograph provided a particularly compelling example of a microbial microenvironment, serving as a protein-rich niche that supported a diverse microbial consortium. Notably, isolates such as *Bacillus altitudinis/pumilus, Oceanobacillus kimchii,* and *Streptomyces rubiginosohelvolus* were identified, all of which thrive in high-salt, low-nutrient conditions, suggesting that albumen prints present unique biochemical properties favorable for microbial colonization. The interaction between microorganisms and the albumen layer, which contains organic compounds such as egg whites and silver salts, may play a significant role in the biodeterioration of photographic materials, potentially leading to discoloration, fading, or other forms of structural damage over time. Other strains isolated from the CHM’s photo book were classified by VITEK, MGs, and ANI as *Microccocus*, *Staphylococcus* and *Kocuria* (Table 1, Fig. 3). These taxa were consistently reported as microbial contamination in historical archives and audiovisual materials (Villalba et al., 2004; Puškárová et al., 2016; Mazzoli et al., 2018; Branysova et al., 2023). They have proven amylolytic, cellulolytic, proteolytic, and/or lipolytic capacities related tobiodeterioration of paper, photographs, and/or other visual material which were mainly determined by culture-dependent methods (Villalba et al., 2004; Branysova et al., 2023). The amylolytic capacities can be used for the consumption of starch from paper (used as a glue in its manufacturing) while the proteolytic activity presents a threat especially to visual materials with a gelatinous binder. Some species with cellulolytic properties pose a risk to collodion visual materials (as the main component of collodion is cellulose nitrate) while others have the ability to produce alkaline serine proteases which are harmful to albumin materials (Villalba et al., 2004; Branysova et al., 2023). Functional genomic analysis of *Streptomyces* sp. CH-036 revealed strong capacities for biodeterioration, including cellulose degradation and pigment biosynthesis, such as isorenieratene, a light-induced pigment. These findings highlight the critical need for targeted conservation measures, particularly avoiding exposure to light and maintaining stable environmental conditions to prevent microbial proliferation. Further experimental validation through controlled humidity exposure tests and protease activity assays would be necessary to confirm this hypothesis and refine conservation strategies for photographic materials.

Notably, *Pseudomonas* strains were exclusively isolated from the photograph, aligning with its role as a frequent contaminant in photographic manufacturing due to its ability to metabolize gelatin and resist silver nitrate (Gadd et al., 1989; Borrego et al., 2010; Kraková et al., 2012b; Sterflinger and Piñar, 2013; Madigan et al., 2014; Grabek-Lejko et al., 2017; Mazzoli et al., 2018; Warner Marien, 2022). Additional research will be required to evaluate the relationship between the *Pseudomonas* isolates of the book and these inorganic compounds.

*Pseudomonas* spp. and *Streptomyces* spp. poses a risk not only because of its potential catalytic properties, but also because of the ability of some members to produce deteriorative pigments. For example, some *Streptomyces* spp. ruined graphic art inside some Etruscan and Roman tombs due to the production of violet pigments, while members of *Pseudomonas* produced both green (in an alkaline environment) and red (in acidic conditions) pigments during wool degradation of historical textiles (Díaz-Herráiz, 2015; Gutarowska et al., 2016; Mazzoli et al., 2018). In accordance with culture-dependent methods, our present genomic analysis over the strain *Streptomyces* sp. CH-036 (Table 2), revealed strong capabilities to degrade components of paper (Fig. 5.D), plus a high number of genes for the biosynthesis of several pigments (Fig. 5.E) including isorenieratene (Fig. 5.F), a light-induced pigment (Takano et al., 2005). These findings should encourage the managers of the museum to take the necessary precautions to avoid exposition of the book to light or favorable conditions for the spreading of the microbe (moist, warm temperatures, etc.).

Some microbes having their optimal growth during the different manufacturing processes of the photo book, they could have been preserved via spores once the manufacturing conditions changed. Indeed, 7 of the 17 strains from the book identified taxonomically by the VITEK method are producers of endospores (*Bacillus* spp.) or resistance spores (*Streptomyces* spp.). Both endospores and resistance spores are metabolically inactive structures, extremely resistant to adverse environmental conditions, such as high concentrations of salts, absence of nutrients or low levels of humidity. The adaptive advantage conferred on bacteria is to allow them to remain as spores for as long as necessary until the conditions are suitable (Madigan et al., 2014). Apparently, this could explain the isolation of the thermophilic strain *Caldibacillus thermoamylovorans* CH-041 which grows optimally at 50°C in the book which is stored at room temperature (20-30°C). The most likely hypothesis is that the thermophile proliferated during the initial manufacturing. Different methods were used to harden the albumen layers, including steaming and storing the albumen-coated papers inside a warm hayloft for half a year. The temperature of a hayloft can sometimes reach 50 °C or more, providing an optimal environment for the growth of CH-041 and other thermophiles like *Geobacillus* reported from a previous albumen photography study (Puškárová et al., 2016). An evolutionary significance of CH-041 is that its genome may have remained unchanged over the past 160 years. A deeper look in to the genomics of CH-041 revealed a scarce amount of genes for antibiotic resistance (Fig. 5.A), and biodeterioration (Fig. 5.D-E), making it the most environment friendly strain reported for the CHM.

### 4.3. Textiles as profilic microenvironments for bacteria proliferation

A particularly intriguing case was observed in Alberdi’s suit, a 19th-century silk and lace garment, where we identified a unique bacterial strain, classified by VITEK as *Actinomyces odontolyticus*. This filamentous bacterium is typically found in human microbiomes, particularly in the oral cavity, and profusely isolated from dental caries (Batty, 1958). Manifestations of infection by *A. odontolyticus* include thoracic, abdominal, pelvic and central nervous system disease (Razok et al., 2022). Recently, it was proposed as driver of Colorectal Cancer (Breau, 2024). Its presence on the textile suggests potential contamination from human handling or storage conditions, a closed cartoon box that avoids contamination and humidity changes. Notably, the possibility that the original owner of the suit, Juan Bautista Alberdi, introduced this bacterium as a child cannot be ruled out, given the high prevalence of dental infections in toddlers and young children. Once established on the fabric, the bacterium’s filamentous morphology may have facilitated its survival by intertwining with textile fibers, contributing to both microbial persistence and the progressive degradation of the material. This hypothesis aligns with scanning electron microscopy (SEM) observations, which revealed extensive colonization of textile fibers by filamentous bacteria (Fig. 2A). The genus *Actinomyces sp*. was previously cited in a list of the most common bacteria responsible for biodeterioration of CH (Grabek-Lejko et al., 2017). A more detailed study of the strain, including genome sequencing should reveal with certainty its taxonomic identity and toxicity, considering the hypothesis that it could have originated from an infected person.

The presence of *Micrococcus luteus* on textiles, a microorganism commonly associated with the accumulation of biological material and potential biodeterioration. Additionally, *Staphylococcus equorum*, found on both textiles and wooden surfaces, is typically linked to environments with human interaction, possibly contributing to the degradation of protein-based materials. Likewise, the *Bacillus* genus, including *Bacillus simplex and Bacillus licheniformis*, was primarily found on wooden structures, suggesting their potential role in the breakdown of cellulose and lignocellulosic compounds.

### 4.4. Furniture and wooden objects associated bacteria

The museum’s wooden furniture subjected to sampling is currently exposed to the air during the year, and suffers the dynamics in temperature and humidity that can enhance bacterial growth. However, they also are affected by frequent procedures of cleaning and disinfection, as well as repairs and preventive actions against wood decay. Frequently, the wooden material in the museum is sensible to other pests, such as insects. For this reason, it is not expected to these wooden items to suffer biodeterioration driven primarily by bacteria. In spite of this, the isolates from the wooden parts of the chair, table, window, washing trough, and main access door could have cellulolytic or lignocellulolytic capabilities required for the degradation of wood. The most commonly reported bacteria associated with wood-decay environments are genera such as *Clostridium*, and cosmopolitan taxa such as *Bacillus* (Liu et al., 2018; Pyzik et al., 2021), both isolated from the aforementioned elements (Table 1). A notable high number of genes related to the breakup of cellulose and hemicellulose were found in the genomes of *Microbacterium* sp. CH-007, *Kocuria* sp. CH-021, and *Microbacterium* sp. CH-015 (Fig. 5.D), which were isolated from the chair wood, the entrance door and the window respectively (Table 1, Fig. 3). This could be supporting some kind metabolic activity over the wooden compounds of the furniture. Moreover, the strongest profiles of heavy metal resistances belong to CH-021 and CH-015, and are probably the reflection of the chemical treatments used over the years to preserve the wood from decay. Most of these wood preservatives recruit heavy metals in their formulas, like copper chromate arsenate (CCA) which was widely used in the past but now restricted due its toxicity (Woźniak, 2022). Finally, SEM micrographs of the washing trough showed the presence of filamentous bacteria probably from the genus *Streptomyces* or related (Fig. 2.D). Previous studies showed that filamentous actinomycetes are associated with wood decay (Antai and Crawford, 1981; Jayasinghe and Parkinson, 2008), while other claim that their presence in wood could prevent worst scenarios of biodeterioration like those caused by fungi (Mortabit et al., 2015; Cai and Kuo, 2022).

One of the strains we isolated from the chair textile was classified as *Bacillus licheniformis*, a species which is list of pathogens infecting humans (Bartlett et al., 2022). *B. licheniformis* was reported in several cases of infection, even in a patient with no history of any immune deficiency (Haydushka et al., 2012). Another intriguing species is *Staphylococcus equorum*, isolated from both the textile chair and the washing trough. *S. equorum* is a commensal and obligate aerobe usually found on the skin of various farm animals, such as horses, dairy cattle and goats (Vermeersch et al., 2023; Reydams et al., 2024; Senoussi et al., 2024). This microorganism probably has colonized the seat and meat washing trough during the nineteenth century, when people of the city interacted routinely with farm animals and used horses as their primary vehicle or cattle as meat source.

### 4.5. Metabolic traits and biotechnological potential of heritage bacteria

The identification of biosynthetic gene clusters (BGCs) related to pigment production and antimicrobial compound synthesis in several isolates carries significant implications for the preservation and study of heritage materials. Pigment biosynthesis pathways—such as those detected in *Streptomyces* sp. CH-036, which includes genes for the light-induced pigment isorenieratene—may contribute to visible discoloration and irreversible staining of culturally significant surfaces. These pigments, often photoreactive and chemically stable, can cause localized chromatic alterations, especially in substrates such as textiles, photographs, and historical paper that are sensitive to light exposure.

In parallel, the detection of BGCs encoding antimicrobial compounds—such as lanthipeptides and ε-poly-L-lysine—suggests that certain strains may exert ecological pressure on their microbial surroundings. These bioactive compounds can inhibit susceptible taxa and promote the survival of resistant microorganisms, potentially influencing the microbial succession and stability on heritage objects. This microbial antagonism may contribute to the reduced diversity observed in specific niches, such as the surfaces of textiles or wooden elements with limited nutrient input.

Together, these metabolic features reveal a dual impact on conservation: while some microbial metabolites contribute directly to biodeterioration through staining or enzymatic degradation, others modulate microbial community dynamics, potentially altering the long-term colonization patterns on heritage materials. These insights underscore the importance of incorporating functional microbial traits—not just taxonomic identification—into the design of informed, preventive conservation strategies.

Moreover, the genomic profiles of the isolated bacteria reveal a promising biotechnological potential beyond their role in biodeterioration. Several *Bacillus* and *Pseudomonas* strains identified in this study have previously been proposed as candidates for bioprecipitation of calcium carbonate crystals, a process with practical applications in biorestoration and consolidation of stone and plaster monuments. Notably, *Bacillus megaterium* and *Bacillus simplex*, both detected among the isolates, are among the most effective carbonate bioprecipitators and are used in protective treatments for historical architecture (Paramo-Aguilera, 2012; Pyzik et al., 2021). In addition, members of the *Bacillus* genus have been suggested for use in biotreatments of textiles and paper artifacts, particularly for their enzymatic capabilities in cleaning and stabilizing cellulosic and proteinaceous materials. Similarly, *Micrococcus* spp. are known for their capacity to produce bioactive molecules and enzymes of interest to the pharmaceutical and bioremediation industries, including antimicrobial compounds and pollutant-degrading enzymes (Díaz-Herráiz, 2015).

Importantly, our genomic analyses revealed that *Streptomyces* sp. CH-036 and *Kocuria* sp. CH-021 harbor genes involved in the synthesis of valuable secondary metabolites and the degradation of xenobiotic compounds (see Fig. 5.F and section 3.4). These findings position these strains as promising sources of novel biotechnological resources and reinforce the concept of cultural heritage environments as overlooked reservoirs of useful microorganisms. Further exploration of these isolates could yield new applications in conservation, biotechnology, and environmental remediation.

## CONCLUDING REMARKS AND PROSPECT

This study provides the first comprehensive microbiological survey of the Casa Histórica de la Independencia Museum, revealing a rich and complex microbial landscape shaped by historical, material, and environmental factors. Through the integration of scanning electron microscopy, microbial isolation, MALDI-TOF MS, and genome sequencing, we uncovered distinct microbial communities inhabiting both outdoor and indoor surfaces of cultural heritage significance.

The comparative analysis between the Outdoor Urban Microbiome (OUM) and the Indoor Museum Microbiome (IMM) revealed striking contrasts. Exterior surfaces, such as the entrance door and walls, were dominated by stress-tolerant and resilient taxa—primarily from the phyla Actinobacteriota and Firmicutes—reflecting their adaptation to fluctuating environmental conditions, UV radiation, and atmospheric pollutants. In contrast, the IMM showed greater taxonomic richness and niche specialization, shaped by more stable temperature, humidity, and limited light exposure. Indoor isolates exhibited strong substrate-specific associations, particularly in organic-rich materials like textiles and albumen-based photographic prints, where human contact and historical use played an important role in microbial transmission.

Artifacts such as the 19th-century albumen print photograph and Alberdi’s silk suit emerged as microecological hotspots, each serving as distinct reservoirs of specialized bacteria. These niches supported extremophilic and potentially novel strains, including *Caldibacillus thermoamylovorans*, *Streptomyces rubiginosohelvolus*, and *Oceanobacillus kimchii*. Their presence reinforces the idea that heritage materials not only face biodeterioration risks but also harbor microbial biodiversity of potential biotechnological value.

Functional genome annotation highlighted a dual role of these microbes: some strains possess genes linked to biodeterioration (e.g., cellulose degradation, pigment biosynthesis), while others exhibit traits with positive applications, such as enzymatic degradation of environmental pollutants. These findings illustrate how cultural heritage microbiomes represent both a conservation challenge and an untapped biological resource.

Understanding microbial colonization patterns at the interface of cultural heritage and environmental microbiology is critical for designing more informed, bio-aware conservation strategies. This includes tailored preservation protocols that address not only visible deterioration but also the metabolic capacities and ecological interactions of microbial communities. Future work should expand to metagenomic and metatranscriptomic approaches, allowing for a deeper exploration of microbial dynamics, functional activity, and long-term succession on heritage substrates.

Ultimately, this study highlights the importance of heritage sites not only as repositories of national memory but also as living ecosystems—microbial archive that reflect centuries of human history, environmental exposure, and material transformation. Protecting this biological dimension of cultural heritage will be key to its sustainable preservation.

## AKNOWLEDGEMENTS

The authors sincerely thank the Museo Casa Histórica de la Independencia staff, especially former Director Cecilia Guerra Orozco and Cecilia Barrionuevo and Ana Oliva from the restoration, conservation and museology department, for kindly granting access to the museum’s collections. All authors are members of the National Research Council (CONICET) in Argentina. VHA was a recipient of a Georg Foster Scholarship for Experienced Researchers, Alexander von Humboldt Foundation (2021-2023) and of a grant of the Williams Foundation (2023-2024). Electron micrographs used in this study were taken at the Center for Electron Microscopy (CIME) belonging to UNT and CONICET, in Tucumán, Argentina.

## AUTHOR CONTRIBUTIONS

VHA had the original project idea, designed and conceptualized the research. DAR and VHA wrote the manuscript. FSG, NA, MCG and VHA performed the sampling. FSG, NA, MCD and MJSM worked in the isolation and characterization of strains and curated the CEBAC collection. MCG performed the photo documentation during the sampling in the museum. FSG, LM, HE performed the electron microscopy analyses. DGAR performed the genomic and bioinformatic analysis. VHA also provided funding, infrastructure, consumables and equipment for the experiments. All authors read and approved the final manuscript, which is available at biorxiv preprint server https://doi.org/10.1101/2025.03.28.645971

